# Tipping-point transition from transient to persistent inflammation in pancreatic islets

**DOI:** 10.1101/2024.03.10.584271

**Authors:** Thomas Holst-Hansen, Pernille Yde, Mogens H. Jensen, Thomas Mandrup-Poulsen, Ala Trusina

## Abstract

Type 2 diabetes (T2D) is associated with a systemic increase in the pro-inflammatory cytokine IL-1β. While transient exposure to low IL-1β concentrations improves insulin secretion and β-cell proliferation in pancreatic islets, prolonged exposure leads to impaired insulin secretion and collective β-cell death. IL-1 is secreted locally by islet-resident macrophages and β-cells; however it is unknown if and how the two opposing modes may emerge at single islet level.

We investigated the duality of IL-1β with a quantitative in-silico model of the IL-1 regulatory network in pancreatic islets. We find that the network can produce either transient or persistent IL-1 responses, when induced by pro-inflammatory and metabolic cues. This suggests that the duality of IL-1 may be regulated at the single islet level. We use two core feedbacks in the IL-1 regulation to explain both modes: First, a fast positive feedback in which IL-1 induces its own production through IL-1R/IKK/NF-κB pathway. Second, a slow negative feedback where NFκ-κB upregulates inhibitors acting at different levels along the IL-1R/IKK/NF-κB pathway – IL-1 receptor antagonist and A20 among others. A transient response ensues when the two feedbacks are balanced. When positive feedback is dominating over the negative islets transit into the persistent inflammation mode. Consistent with several observations, where the size of islets was implicated in its inflammatory state, we find that large islets and islets with high density of IL-1β amplifying cells are more prone to transit into persistent IL-1β mode.

Our results are likely not limited to IL-1β but general for the combined effect of multiple pro-inflammatory cytokines and chemokines. Generalizing complex regulations in terms of two feedbacks of opposing nature and acting on different time scales provides a number of testable predictions, which call for dynamic monitoring of pro-inflammatory cytokines at the single islet level.

**AUTHOR SUMMARY:** Different expression or activity dynamics of the same proteins or signaling molecules can lead to opposing fates in cells and tissues. While it is known that brief and prolonged exposure to pro-inflammatory cytokine IL-1β have opposing effects on the functionality and viability of pancreatic β-cells, it is unclear if and how these differences in dynamics may arise at the single islet level. We use a mathematical model of the core feedback loops in the IL-1β regulatory network to show that transient and persistent responses are the two characteristic dynamic modes of the IL-1β response. The likelihood of each mode depends on systemic inflammation and elevated glycaemia and free fatty acids levels. We find that large islets are more prone to transit into the persistent mode, which may provide an explanation for why large islets are underrepresented in type 2 diabetes patients.

## INTRODUCTION

When cells of higher eukaryotes respond to perturbations in cellular homeostasis, they often choose between two primary strategies: They either adapt and repair the inflicted damage or commit to programmed cell death. The final decision depends on the severity of damage as reflected in the intensity of activation of signaling pathways and levels of regulatory proteins. It becomes increasingly clear that the same proteins that act protectively under remediable stresses can induce cell death when cells face severe or prolonged stress. Furthermore, the switch from adaptive to detrimental action is often sudden and irreversible [1, 2]. The examples of life/death duality span across kinases (Ire1 [3], PERK [4, 5], JNK [6]), transcription factors (p53, NF-κB [7, 8]), DNA repair proteins (BRCA1 [9]), mitochondrial factors (Bax and Bak [10]), ROS signaling molecules (H_2_O_2_ [11, 12]) and cytokines (IL-1[13]).

While the intracellular components of these life/death signals are involved in single cell decisions, diffusible or secreted factors (e.g. H_2_O_2_ and IL-1) are used in intercellular communication and influence collective decisions. There are several exciting findings uncovering the dynamical nature of the life/death transitions in single cells [1, 2], however the understanding of protein duality at the tissue level remains an uncharted area. It is unclear how the geometry of the tissue and inflicted damage influence the collective decision.

We address these questions with the help of theoretical and in-silico modeling of IL-1β regulation in islets of Langerhans. The dual mode of action of IL-1β cytokine on the pancreatic β cell is well established: while transient exposure to low IL-1β concentrations improves insulin secretion and promotes β-cell survival, prolonged exposure to high levels of IL-1β, as in chronic inflammation, leads to impaired insulin secretion and β-cell death [14-17]. The islets of Langerhans present a unique model system for quantitative understanding, as islet geometry and their cellular composition is well described across many species [18-20], the IL-1β regulatory network is well studied, and islets comprise a computationally amenable number of IL-1β secreting cells, ISCs.

Furthermore, IL-1β plays an important role in the progression of type 1 and 2 diabetes [13, 15]. Type 2 diabetes (T2D) results when pancreatic β-cells fail to compensate for increased insulin needs as a consequence of insulin resistance. T2D is associated with systemic low-grade chronic inflammation and immune-cell infiltration of insulin-sensitive tissues and pancreatic islets. Proof of concept of the role of IL-1 in the progression of T2D, stems from clinical trials, where attenuating IL-1 signaling improves glycaemia by enhancing β-cell function [21]. Interestingly, both infiltrating macrophages and pancreatic β-cells are believed to be sources of IL-1β in pancreatic islets [16]. IL-1β synthesis is tightly regulated and requires two signals (Figure 1A). A priming signal (Signal 1), in the form of pathogen associated molecular patterns (PAMPS) and pro-inflammatory cytokines including IL-1β itself activates NF-κB, which in turn induces expression of the IL-1β precursor pro-IL1β. This is an inactive form of IL-1β that remains inside the cell. A second signal (Signal 2), represented by high concentrations of glucose and free fatty acids (FFA) and Damage Associated Molecular Patterns (DAMPS), activates the NLRP3 inflammasome for pro-IL-1β cleavage by caspase-1. This converts pro-IL-1β into its mature form, which is exported into the extracellular space [15]. Exposure of β-cells to both Signal 1 and Signal 2 results in an amplifying feedback loop in which IL-1β induces its own expression. As a result, the β-cells may contribute to chronically elevated IL-1β levels in a self-sustaining manner establishing a vicious cycle [13, 14, 22]. If this auto-stimulation is not balanced it may result in chronic inflammation and cause islet pathology and eventual death. Furthermore, IL-1β acts as chemoattractant for immune cells such as macrophages. The attracted macrophages may further contribute to the chronic levels of IL-1β in the inflamed islets as the macrophages use the same set of cues as β-cells to upregulate mature IL-1β.

**Figure 1.**
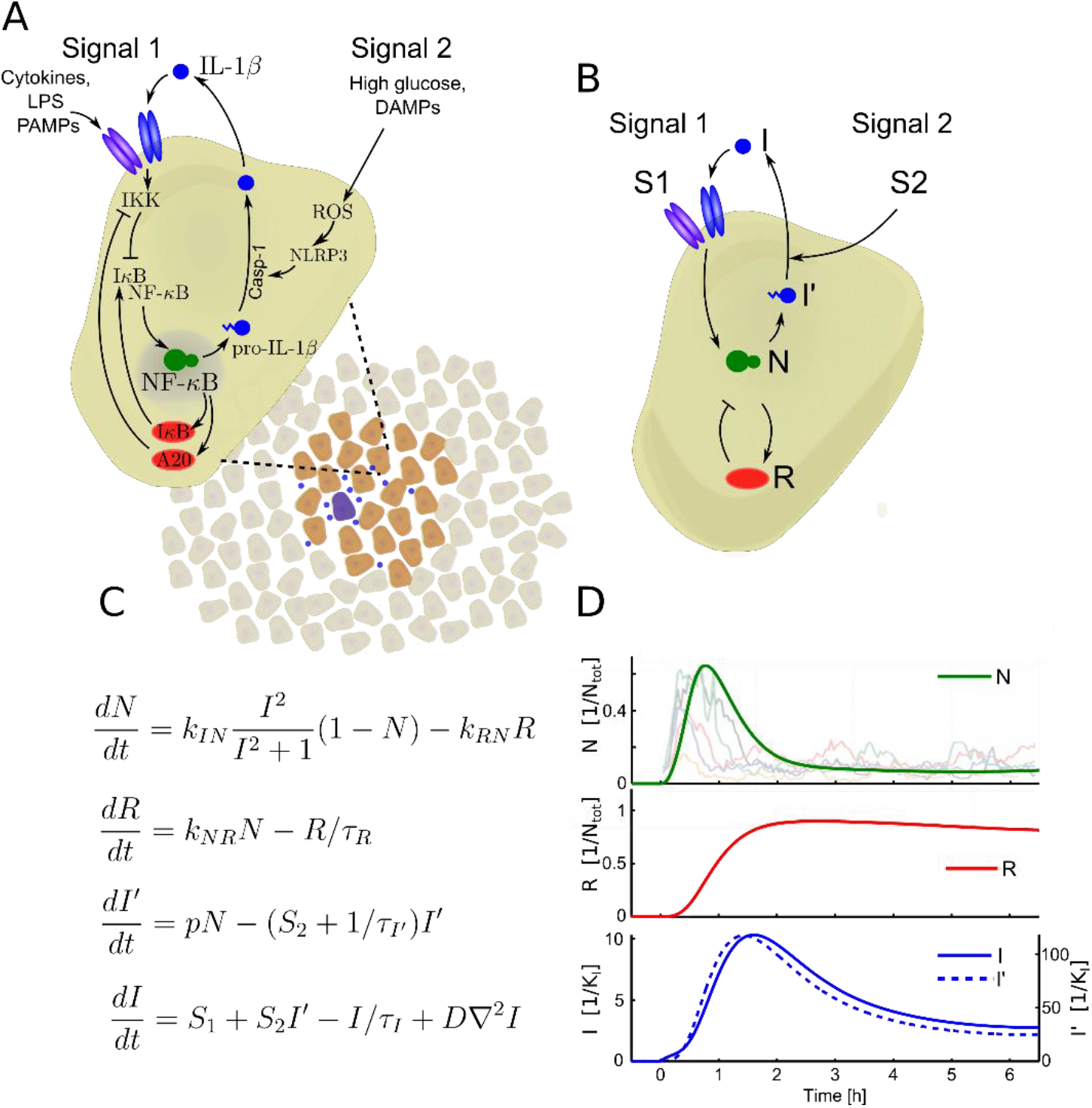
Schematic of the IL-1β regulatory loops in a pancreatic islet and the corresponding in-silico model.. **A** The main signaling pathways of IL-1β in a β-cell or immune cell infiltrating the islets. The regulatory system can be grouped into two pathways; Signal 1 leads to NF-κB activation and production of pro-IL-1β. Signal 2 activates the NLRP3-inflammasome and leads to subsequent caspase-1 activation that cleaves pro-IL-1β into mature IL-1β. Together Signal 1 and 2 compose a self-amplifying feedback loop by which IL-1β stimulates its own production. We model the islet of Langerhans in two dimensions as a cluster of IL-1 secreting cells, surrounded by non-secreting cells, which do not respond to IL-1β. In the model a random set of cells act as initial sources of Signal 1. **B** Simplified model of the IL-1β signaling pathways in a single cell. The model includes four variables representing pro-IL-1β (*I’*), IL-1β (*I*), NF-κB (*N*) and proteins regulating NF-κB through negative feedback loops (grouped into the variable *R*). There are two free parameters in the model: *S*_*1*_ represents stimuli by a cytokine source (e.g. IL-1β) and is related to Signal 1 and *S*_*2*_ is the maturation rate of IL-1β that is related to caspase-1 activity and hence Signal 2. **C** Model equations describing IL-1β dynamics. **D** The output from a single IL-1 secreting cell is fitted to NF-κB data such that the typical NF-κB response peaks within ≈ 1 hr of stimulus, overlay is from [30]. Here we show the response of a single cell stimulated by a persistent *S*_*1*_ = 5.0 hr^-1^ and *S*_*2*_ = 0.5 hr^-1^.

In spite of the potential importance of IL-1β in T2D progression and the extensive knowledge on IL-1β regulation in pancreatic β- and immune cells, the nature of IL-1β transition from an adaptive to a deleterious state remains largely unknown. It is not known, if the transition in single islets is gradual, with a gradually increasing fraction of dysfunctional cells, or if it is a collective decision, with all cells transitioning from a functional to dysfunctional state simultaneously. Remarkably, islet geometry correlates with sensitivity towards inflammation: the immune cells infiltrate *large* islets first [15]; large islets are underrepresented in T2D patients [23] and knocking down NLRP3 inflammasome in murine models of T2D caused a significant increase in islet sizes compared with wildtype obese mice [24]. To date the mechanisms behind the correlations between islet geometry and its propensity to inflammation are unclear.

With the help of the in-silico model, we find that when islets are exposed to pro-inflammatory and nutritional cues, there are two characteristic modes of IL-1 upregulation: *transient* or *persistent*. More importantly, the model predicts that while the increase in number of pathological islets exposed to pro-inflammatory and nutritional cues may be slow and gradual, the transitions to the persistent mode in single islets are rapid and sudden. Our analysis shows that the transition is sensitive to islet architecture and size: while larger islets with dense cores of ISC are more prone to chronic inflammation, the small sized and folded organization of human islets adds to the tight regulation of IL-1β and offsets the onset of the persistent mode.

## RESULTS

### Mapping known IL-1 regulations to a mathematical model

In order to minimize the number of unknown parameters and variables, we aimed to construct a simple model, which captures the qualitative behavior of the biological system. In our earlier theoretical studies, we have shown that cytokine auto-stimulation through the NF-κB pathway belongs to a class of phenomena known as excitable media [25, 26]. Here we modify an earlier model [25] to describe the NF-κB mediated inflammatory response in pancreatic islets. It has been shown that, in addition to macrophages, β-cells contribute to IL-1β -accumulation in the islets [27]. In the current model we do not differentiate between the two, and refer to them as IL-1β Secreting Cells (ISC). We assume that other endocrine cells types (α, δ and γ) do not contribute to cytokine secretion.

The cells were arranged on a hexagonal grid and their positions were fixed. IL-1β was assumed to diffuse freely between cells; based on a molecular weight of 17 kDa the diffusion coefficient of freely diffusing IL-1β is estimated to be D = 20 µm^2^/s [28]. While we do not explicitly model small blood vessels penetrating into islets, we account for the fact that β-cells are typically in close vicinity to the blood, by using plasma clearance rates of IL-1β as an estimate of the effective IL-1β half-life in the islets (*τ*_*I*_ = 0.17 *hr*) [29].

### Model of IL-1β regulation in an ISC

We reduced the detailed regulatory network of IL-1β shown in Figure 1A, to an effective regulatory network (Figure 1B). The model includes four dynamic variables: *I, I’, N* and *R. I* represents the concentration of IL-1β, *I’* is the concentration of pro-IL-1β, *N* represents the concentration of nuclear NF-κB (i.e. active in transcription) and R represents the combined effect of regulating proteins which inhibit active NF-κB (e.g. IκBα, A20, cylindromatosis (CYLD), etc.). While the IκBα negative feedback is significantly faster, compared to A20 and CYLD, combining them together in one slow feedback produces qualitatively the same results as long as one does not aim to reproduce the lower amplitude secondary oscillations [25].

The dynamics of each variable is described by an ordinary differential equation:

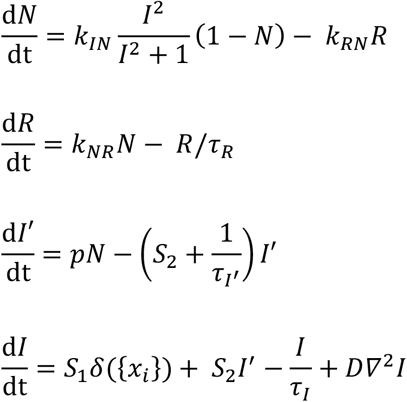

The parameters *k*_*IN*_, *k*_*RN*_, *k*_*NR*_ and *τ*_*R*_ have been fixed by fitting the model to experimental data for TNF-induced NF-κB dynamics in single cells [30], see Figure 1D. Notice that only IL-1β diffuses in the islet extracellular space whereas the other proteins are intracellular. While the data on IL-1β induced NF-κB responses in single cells is not available; population averaged data supports transient activation of NF-κB with a peak around 2 hours [31]. Further details on model parameters and approximations are presented in detail in the Methods section.

Our investigation focuses on the parameters *S*_*1*_ and *S*_*2*_, related to the levels of initiating inflammatory (Signal 1) and metabolic cues (Signal 2), and how these parameters affect the dynamics of the model.

### Signal 1: Inflammatory cues

Although IL-1β could be produced solely by islet-invading immune cells such as macrophages, it has been hypothesized that the initial increase of IL-1β is mainly produced by stressed β-cells. This increase may subsequently attract immune cells, adding further to the increasing levels of IL-1β [14]. In our model, we do not specify the originators of the initial increase of IL-1β, but simulate Signal 1 by adding a basal production of IL-1β to a number of randomly chosen ISCs. These sources of Signal 1 act as initiators of the model, allowing us to study how the ICSs respond and regulate IL-1β when exposed to an external stimulus. Signal 1 is thus characterized by both the strength of basal IL-1β production, *S*_*1*_, and the number of sources, *ns*. We stress that the initial sources described by the *S*_*1*_ parameter reflect a cumulative effect of other NF-κB activating agents such as TNF, TLR, PAMPs or other cytokines, but for the purpose of our model, we describe these as IL-1β sources in order to minimize the number of variables. While we choose to model *S*_*1*_ as a few discrete sources, we have tested that the model results hold if instead *S*_*1*_ is equally distributed across all cells. This configuration is in particular relevant for cases where Signal 1 represents low-grade inflammation stemming from sources outside the islets, with low levels of pro-inflammatory cytokines circulating in the blood [32].

### Signal 2: Nutritional cues

The severity of T2D is positively correlated with the levels of glucose, the amount of free fatty lipids concentration in the blood and DAMPs in the extracellular space. These factors all contribute to the activity of the NLRP3 inflammasome and in turn caspase-1, see Figure 1A. In order to construct a simple model, we merge the effects which contribute to an increased caspase-1 activity into the *S*_*2*_ parameter (see Figure 1C). But since caspase-1 actively cleaves pro-IL-1β into mature IL-1β, the closest biological interpretation of *S*_*2*_ is the concentration of caspase-1. *S*_*2*_ in combination with the production rate of pro-IL-1β, *p*, constitutes the strength of the positive feedback by which β-cells amplify the local IL-1β concentration. This becomes a crucial parameter in the qualitative behavior of the simulated islets.

The ISC synthesize IL-1β and secrete it into the extracellular space, where IL-1β acts in an autocrine and paracrine manner. In each simulation, islets are initiated in a steady-state of low activity with no stimulation, i.e. NF-κB is inactive and *S*_*1*_ equals zero for all ISCs. At time zero the islets are stimulated by a number of small inflammation sources – technically *S*_*1*_ is set equal to a non-zero value for a discrete set of spatial positions. The sources are only non-zero for a finite time, after which the *S*_*1*_ parameter is set equal to zero, and we monitor the islet response.

## Model validation

### The model predicts transient and persistent production of IL-1β in single islets, in correspondence with the IL-1β dual effect

The bimodal effects of IL-1 on β-cell function and viability are associated not only with the concentration of IL-1 but also with the duration of exposure. Transient exposure leads to protective effects while prolonged exposure promotes cell death [14, 33]. As pancreatic islets have been shown to mount a significant local IL-1 response, among the endocrine islet cells and infiltrating macrophages, we wanted to investigate if the two distinct temporal profiles of IL-1 may emerge on the single islet level. To investigate this, we monitored how levels and duration of the IL-1 production depend on nutritional (*S*_*1*_) and inflammatory cues (*S*_*2*_) within an in-silico islet.

We found that whereas low levels of *S*_*1*_ and *S*_*2*_ result in *transient* upregulation of IL-1β, high levels of either *S*_*1*_ or *S*_*2*_ (or both) will result in *sustained high* levels of IL-1β (Figure 2A-B). When an islet with a low *S*_*2*_ is stimulated by a small number of inflammatory sources (low *S*_*1*_), it responds with a pulsating NF-κB activity and a similar pulsating amplification of IL-1β (Figure 2A). Once the sources are removed, the islet stops pulsing and returns to the resting state (see supplemental movies S1 and S4). Hence, the islet exhibits a transient response to the transient inflammatory cues. If a similar islet is exposed to either higher Signal 1 (more initial sources of inflammation) or to higher Signal 2 (e.g. higher caspase-1 activity) the cells will collectively amplify the IL-1β concentration and transition into a “locked” state of constantly elevated IL-1β (see movies S2, S3, S5 and S6 and Sec. II in SI). In this state the islet will sustain the high levels of IL-1β, and will not settle back to the resting state - even when the initial IL-1β sources are removed. The locked state simulates a state of chronic inflammation, where prolonged exposure to high levels of IL-1β has been reported to have deleterious effects leading to impaired insulin secretion and β-cell death [14-17]. Similarly to our earlier work, [25], dynamical systems analysis showed that the locked state is a consequence of an overactive positive feedback in ISC, which prevents the fast variables (I, I’ and N) from undergoing a saddle node bifurcation and return to low cytokine levels. These results are consistent with the dual role of IL-1β reported in diabetic patients (or lab animals), where it can both facilitate cell survival, the transient response, or enter a state of chronic inflammation, the locked state [14, 33]. Notably, the islets display at least one NF-κB/IL-1β pulse in response to a stimulus, and therefore even the islets which eventually lock also display an initial transient IL-1β response (Figure 2A-B), suggesting that the chronic inflammation state will always follow after the initial acute phase with transient IL-1β.

**Figure 2.**
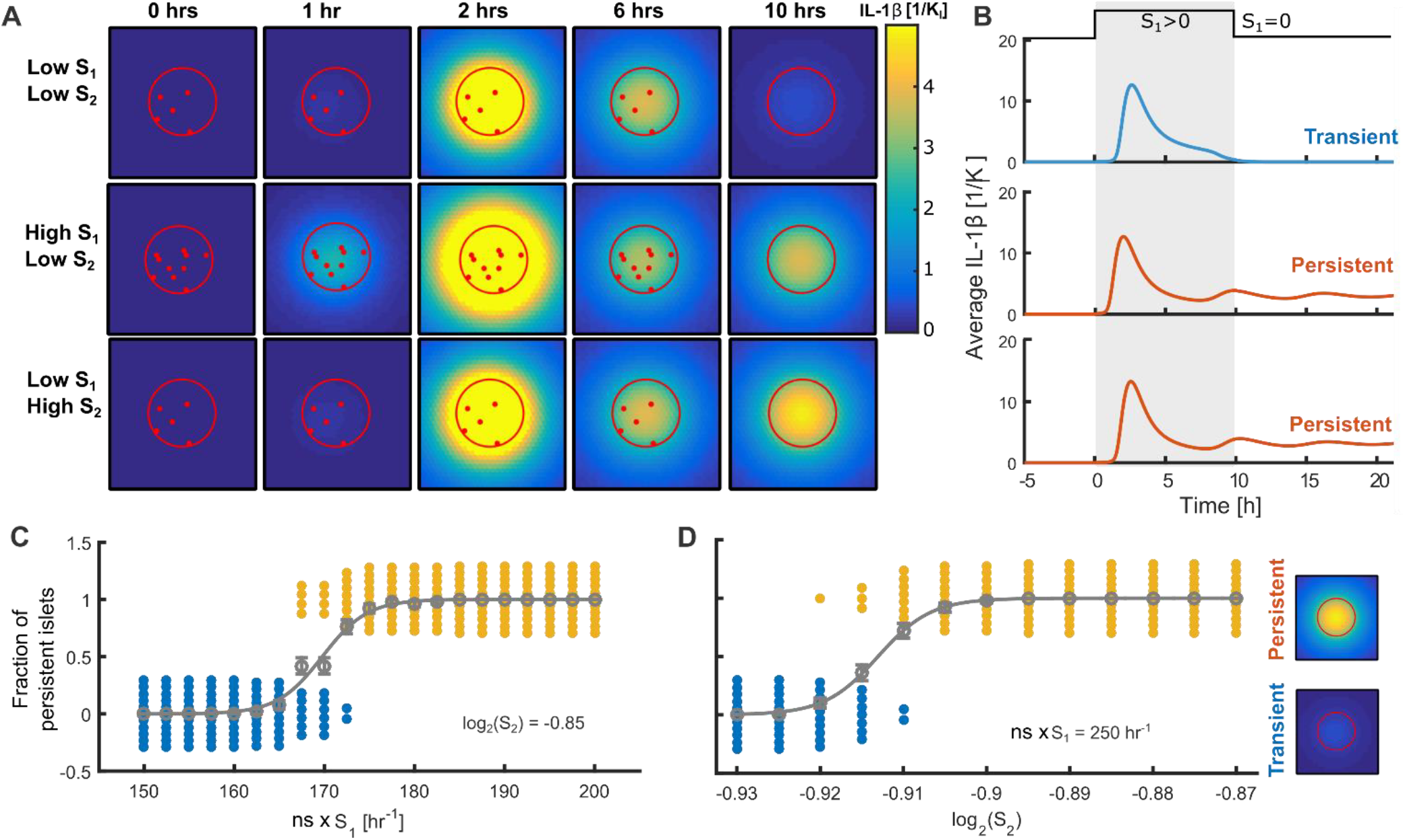
Transient and persistent IL-1 modes emerge in single islets. **A**. Snapshots of IL-1 in islets subject to different combinations of high and low *S*_*1*_ and *S*_*2*,_ stimulated with *S*_*1*_ for ten hours. ISCs are located within the red circle and non-secreting cells outside. The sources of *S*_*1*_ are shown as red dots. **B**. Time courses of IL-1β averaged over all cells in the islet corresponding to the islets in **A**. Under all conditions, islets display an initial strong secretion of IL-1β. At low *S*_*1*_ and *S*_*2*_ the islets respond transiently and will return to the resting state (Movie S1 in SI). If either *S*_*2*_ or *S*_*1*_ is high, islets can enter a “persistent” state with sustained IL-1β expression (Movies S2 and S3 in SI). **C** The fraction of persistent islets increases with the total *S*_*1*_ dose (number of sources times the *S*_*1*_) **D** The fraction of persistent islets increases with *S*_*2*_. Grey points and error bars show average number of persistent islets, grey line is a sigmoidal fit used to find EC_50_ (see Methods and SI). Overlaid in blue and yellow are the bee-swarm plots, where each dot represents five islets. For both parameters, there is a region of coexistence between persistent islets and transient islets that return to the resting state. The state of individual islets depends on the positions of *S*_*1*_ sources. Here each islet has a radius of 7 ISC. Results for larger sizes are shown in Figure 5.

Having established two distinct modes of IL-1β response we investigated if the transition between the two modes is gradual, with e.g. islets gradually increasing production levels of IL-1β with increasing *S*_*1,2*_ or sudden, with a discontinuous jump from low to high IL-1β with just a small variation in *S*_*1,2*_.

We found that the transition is sharp (Figure 2C-D and confirmed by the larger parameter-scan shown in Figure 3 E-H): A small increase in either Signal 1 or Signal 2 (or both) can have a drastic effect on the probability of locking. Notably, close to the transition, locked and non-locked islets can coexist: despite being exposed to the same *S*_*1*_ and *S*_*2*_ islets lock into a state with sustained level of IL-1β, whereas others show transient behavior. In this regime the fate of an individual islet is determined by the exact geometrical positioning (and timing) of the *S*_*1*_ sources. This finding is in line with the current knowledge that in the same pancreas there can be co-existing subpopulations of inflamed and non-inflamed islets [15] (measured by number of infiltrating macrophages). Our results also hold under the conditions of systemic *S*_*1*_, representing elevated IL-1 in the blood, e.g. secreted by adipose tissue (SI, Sec. I.d and Figure S2).

**Figure 3.**
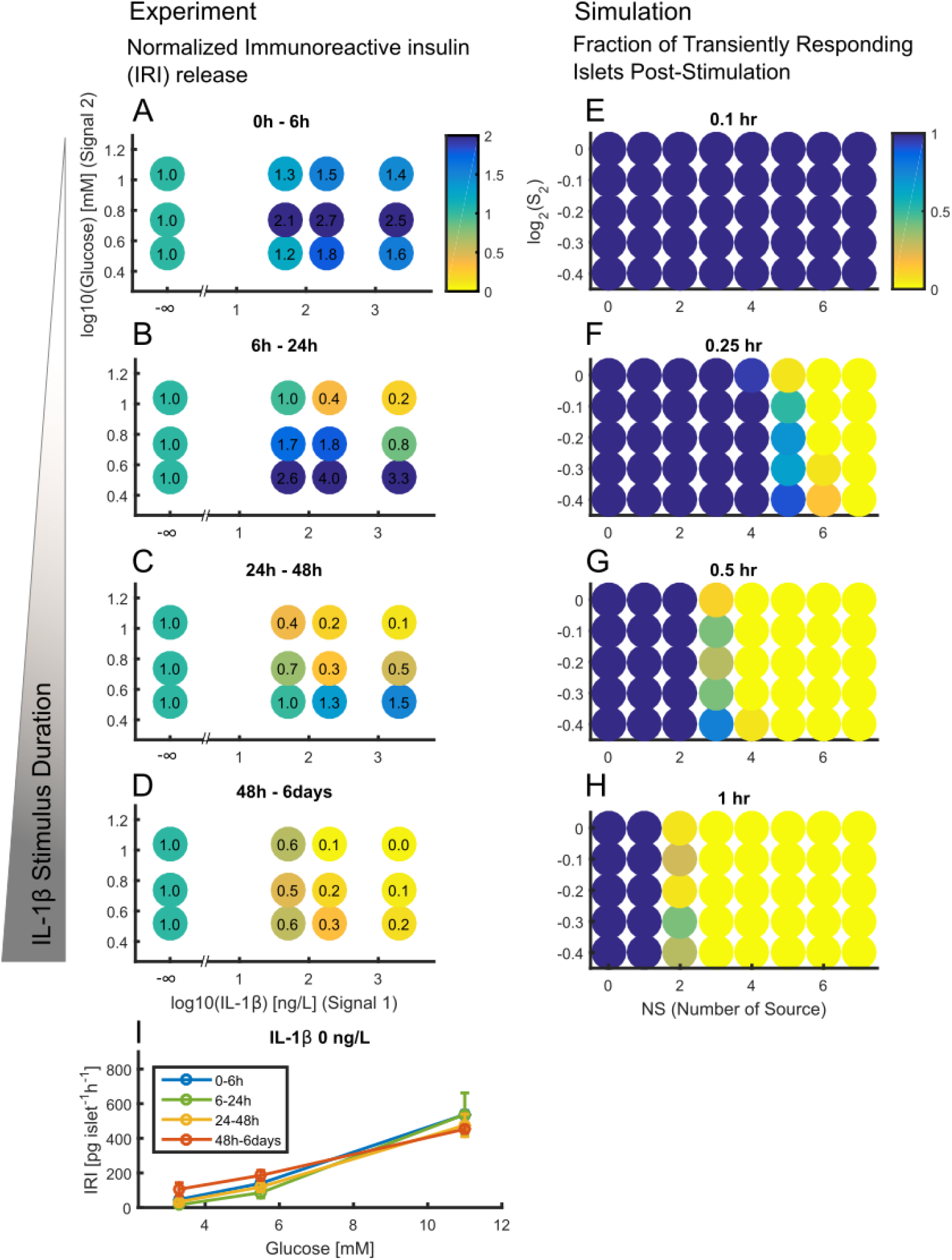
Transition from IL-1 protective to deleterious effects is accelerated under higher *S*_*2*_. **A***-***D** The time course of the Immunoreactive Insulin Release (IRIR) normalized with respect to IRIR in absence of IL-1β. **I** IRIR in absence of IL-1β data from [34]. The IRIR was measured in vitro in islets exposed to different concentrations of glucose and IL-1β. The islets were exposed to increasing durations of IL-1β from **A** to **D**.**F-H** Simulation results showing the fraction of islets responding transiently as function of *S*_*1*_ and *S*_*2*_. Each point consists of 20 islets of a radius of 6.7 cells with randomly placed external sources within the islet. Similar to the experiments, the probability of transitioning to the “locked state” increases as function of external stimulus, inflammasome activation and the duration of external stimulus. Each external source has an *S*_*1*_-value of 25 hr^-1^ and p = 1400 hr^-1^.

### The model recapitulates the fact that glucose narrows the time-window in which IL-1 has beneficial effects

Having established the emergence of transient and persistent modes of IL-1 production within the single islet, we wanted to further test if our model is consistent with the experimental observation that the time window of the beneficial effect from IL-1 is shorter at higher glucose levels (higher *S*_*2*_) [34].

Palmer et. al. [34] measured the insulin response to different glucose concentrations in islets, while islets were exposed to concentrations of IL-1β for increasing durations of time.. To focus on the effects of IL-1β, we have plotted the values reported in [34] in Figure 3A-D, where we normalized insulin responses at each glucose and IL-1β level by the corresponding values in the absence of IL-1β (Figure 3I). Qualitatively, the effects of IL-1 are glucose-independent at brief (0-6 hours, Figure 3A) and long (2-6 days, Figure 3D) exposures. At all glucose levels, brief exposure increases and long exposure inhibits insulin release. The switch between the two modes, which happens at intermediate durations, is however glucose dependent, occurring already after 6 hours at the highest glucose concentration (Figure 3B).

For our model to be consistent with these results, *brief* exposure to *S*_*1*_ should result in a majority of islets in transient mode. *Long exposure to S*_*1*_, on the other hand, should result in a majority of islets in persistent mode. These two cases should be *S*_*2*_-independent, that is the majority of islets will be in the same state at all values of *S*_*2*_. At the intermediate durations, switching from the transient to the persistent mode should happen first where *S*_*2*_ is highest. We find that this is indeed the case (Figure 3 E-H). Notably, the islets exposed to IL-1β for a longer time display an increasingly impaired response to glucose stimulation. This is consistent with our simulations, where the parameter range of *S*_*1*_ and *S*_*2*_ that leads to locked islets, increases with longer exposure to *S*_*1*_.

To our knowledge, it is not known why the switching from insulin-enhancing to insulin-inhibiting modes happens earlier at higher glucose concentrations. While it has been shown that glucose potentiates IL-1β-induced nitric oxide production – mediator of the IL-1β cytotoxic effect, the time dependence has not been addressed[35]. Our results suggest that the time-dependence may come as a result of the modulation of the positive feedback-loop by *S*_*2*_, in that stronger positive feedback accelerates the transition to “locked” state and thus to insulin-inhibiting mode of IL-1β. When we compare simulation and experimental results, the region of parameters where we find locked islets increases much faster in simulations than the corresponding region for the islets with impaired Immunoreactive Insulin Release (IRIR) in the experiments, Figure 3. In the experimental settings, inflammasome activation (signal 2) is likely to increase gradually over time, since the islets are exposed to the combined stress of high glucose and IL-1β. This is not explicitly incorporated in our model; *S*_*2*_ is set to a certain constant value at the beginning of our simulations, but could be made a function of exposure time to fit the experimental transitions.

### Robustness of the results to parameter choices

Islets of Langerhans are subject to cell heterogeneity – not only in different cell types, but also variations in protein levels inside and outside the cells.. The only source of stochasticity in our model has been the position of sources within the islets. Noise in parameters such as inflammasome activation or clearance of inhibitor can have large impact on the local dynamics, however we do not expect this to change the main outcomes of the model

The diffusion constant used in the main results was based on the estimate of diffusion of IL-1β from its molecular weight in cell-free medium; however, binding of IL-1 in the extracellular matrix is likely to result in a lower effective diffusion [36]. We accounted for this by checking that our results do not change qualitatively if the diffusion constant is decreased up to 100-fold, see supplementary movies S7,8,9. Lowering the diffusion constant shifts and decreases the relevant parameter regimes (*S*_*1*_, *S*_*2*_ and p), since less cytokine is removed from the islets. This scenario can lead to interesting phenomena as seen in excitable media such as travelling waves or spiral patterns (see SI Movie S10,11). In rare instances, source position can generate waves that travel around central arteries as in the human and donut islet.

### The effect of IL-1 receptor antagonist on islet fate

Because IL-1β is an established part of T2D pathogenesis, different types of treatments targeting the cytokine have previously been investigated. Clinical studies with anakinra, a recombinant IL-1 receptor antagonist (IL-1Ra) have showed improvements in glycemia and β-cell function [37]. Remarkably, a 3-month anakinra treatment had a durable effect, lasting for over 39 weeks after anakinra withdrawal. Other studies targeted the increased level of cytokines through IL-1β antibodies, with a similar improvement of β-cell function and lowered inflammation [38]. While the mechanism behind the peristent effects after drug withdrawal is unknown, it has been hypothesized that it could result from interrupting the auto-inflammatory positive feedback loops [39]. We set out to test this hypothesis with our model. The result of blocking IL-receptors with the antagonist effectively corresponds to increasing the activation threshold of NF-κB in the model. This can be done in two ways: preventive treatment, where the drug is given before islets have transitioned into the locked state or reactive treatment, where the drug is given after the islets transitioned into the locked state.

We find that preventive treatment offsets the transition by shifting the locked state to higher *S*_*2*_, Figure 4A. in other words islets can withstand higher levels of glycaemia or free fatty acids in the presence of the antagonist. To asses the degree of dose-dependency of the simulated treatment, we have normalized the results in Figure 4A by the untreated results (corresponding to 1.0 K_I_),, see Figure 4C. The normalized values can be thought of as islets “rescued” by the treatment. We find that the effect is dose-dependent in several ways: first, within a certain range of *S*_*2*_ (log_2_S_2_=-0.92 : -0.9) the fraction of “rescued” islets increases with increasing dose. Second, the range of *S*_*2*_ where the treatment has an effect also increases with the dose. While the trend is the same for the case of reactive treatment, where antagonist is added after islets have already locked, we find that reactive treamtnet is less efficient. Rescue of about 50% of the islets required 10-fold higher doses (higher activation threshold) compared with the preventive treatment.

**Figure 4:**
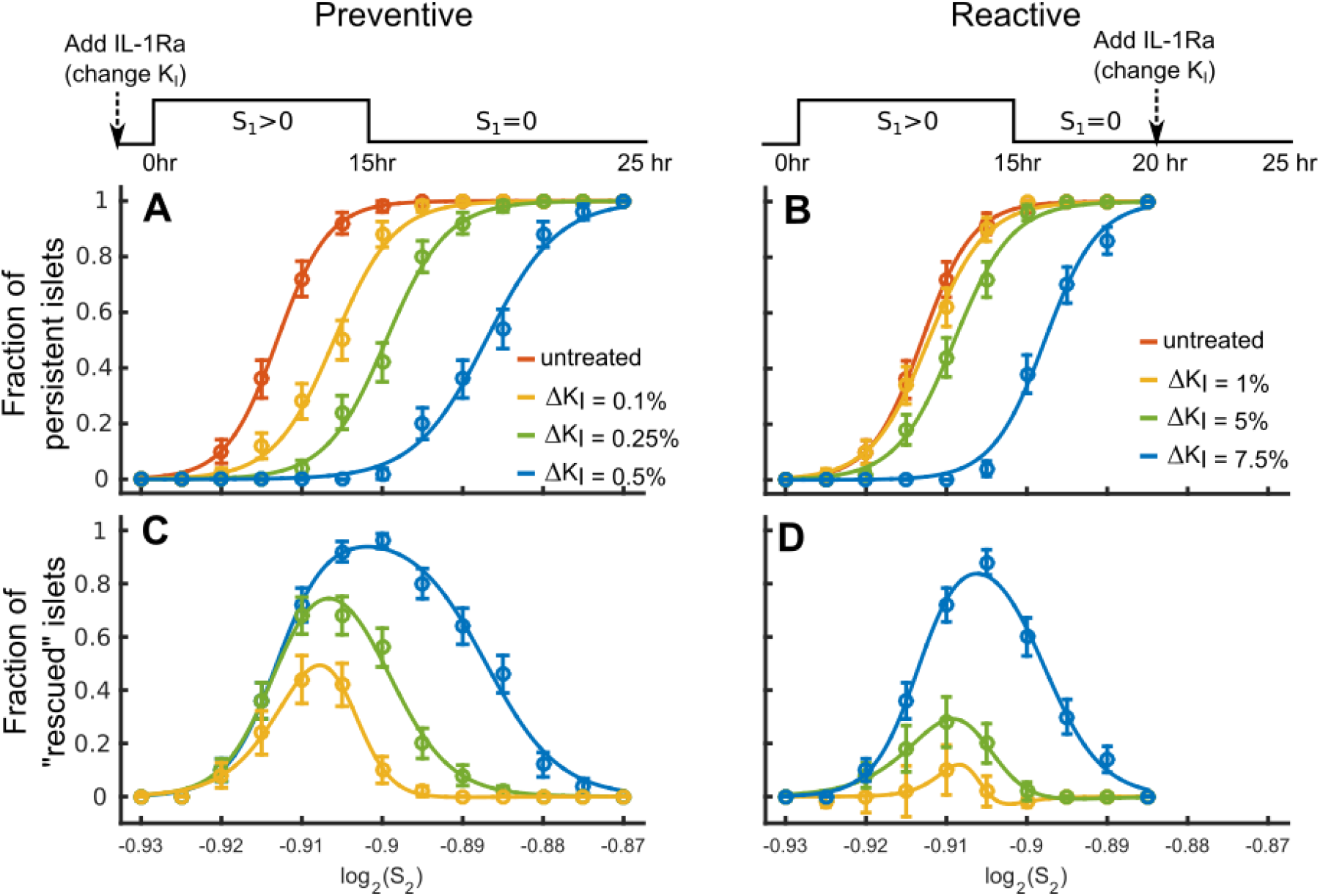
Simulated effect of preventive and reactive treatment with IL-1β antagonists. **A** Preventive treatment is simulated by changing the activation threshold, K_I_, prior to *S*_*1*_ exposure. Increasing K_I_ offsets EC50 to higher *S*_*2*_. Simulations of the untreated case (1K_I_, red). K_I_ is increased by ΔK_I_ 0.1% (yellow), 0.25% (green) and 0.5% (blue). **B** Reactive treatment is simulated by changing activation threshold after *S*_*1*_ exposure. The results of the simulations in the untreated case with (1K_I_, red) and increased activation threshold by 1% (yellow), 5% (green) and 7.5% (blue). **C, D**, The fraction of “rescued” islets represented by the relative change (treatment – no treatment) in fraction of locked islets. The reactive treatment has a weaker effect than preventive treatment, and the range of S_2_ where the treatment has the maximum effect is dose-dependent. All data was fitted by sigmoidal curves (colored lines), see Methods and SI Analysis section. Results are from simulations in islets of radius 6.7 cells.

While interfering with the positive feedback loop through both preventive and reactive treamtents can rescue the islets from locking, we find that removing the treatment (resetting the K_I_ back to pre-treated values) would not have long-lasting effects as long as *S*_*1*_ and *S*_*2*_ are unchanged. It is however known, that the IL-1R antagonists improve glycaemia and FFA levels in T2D patients [39]. This corresponds to lowering *S*_*2*_ prior to treament withdrawal in our model and in this case the effect of single islet unlocking with IL-1RA would be long-lasting.

### Islet shape, size and internal geometry play crucial roles in the fate of islets

Human islets are diverse in size and geometrical arrangement of β-cells. Small islets tend to have a core-mantle shaped structure with a homogenous core of β-cells, surrounded by α-, β- and δ cells. Larger islets are more complex, as the core-mantle geometry is replaced by heterogeneous mixture of α-, β- and δ cells and fenestrated capillaries[23]. Notably, it is large islets that are the first to be lost in T2D patients [23]. While the mechanism is currently unknown, there is strong evidence that inflammation plays an important role. Firstly, Ehses et al. [15] found that in murine models of T2D, macrophages infiltrate large islets first. Secondly, Youm et al. [24] showed that ablation of NLRP3 inflammasome in chronically obese mice protected large islets from inflammation-induced death.

These observations together with our findings suggest that the larger islets may have a higher propensity to lock, and as a consequence would be compromised at lower levels of *S*_*2*_. In order to test this, we first varied the size of the core-mantle shaped islets and monitored the fraction of persistently responding islets with increasing *S*_*2*_ as described in Figure 2D. We found that the transition from transient to persistent IL-1 response indeed occurs at lower values *S*_*2*_ when islets are larger (Figure 5A).

**Figure 5:**
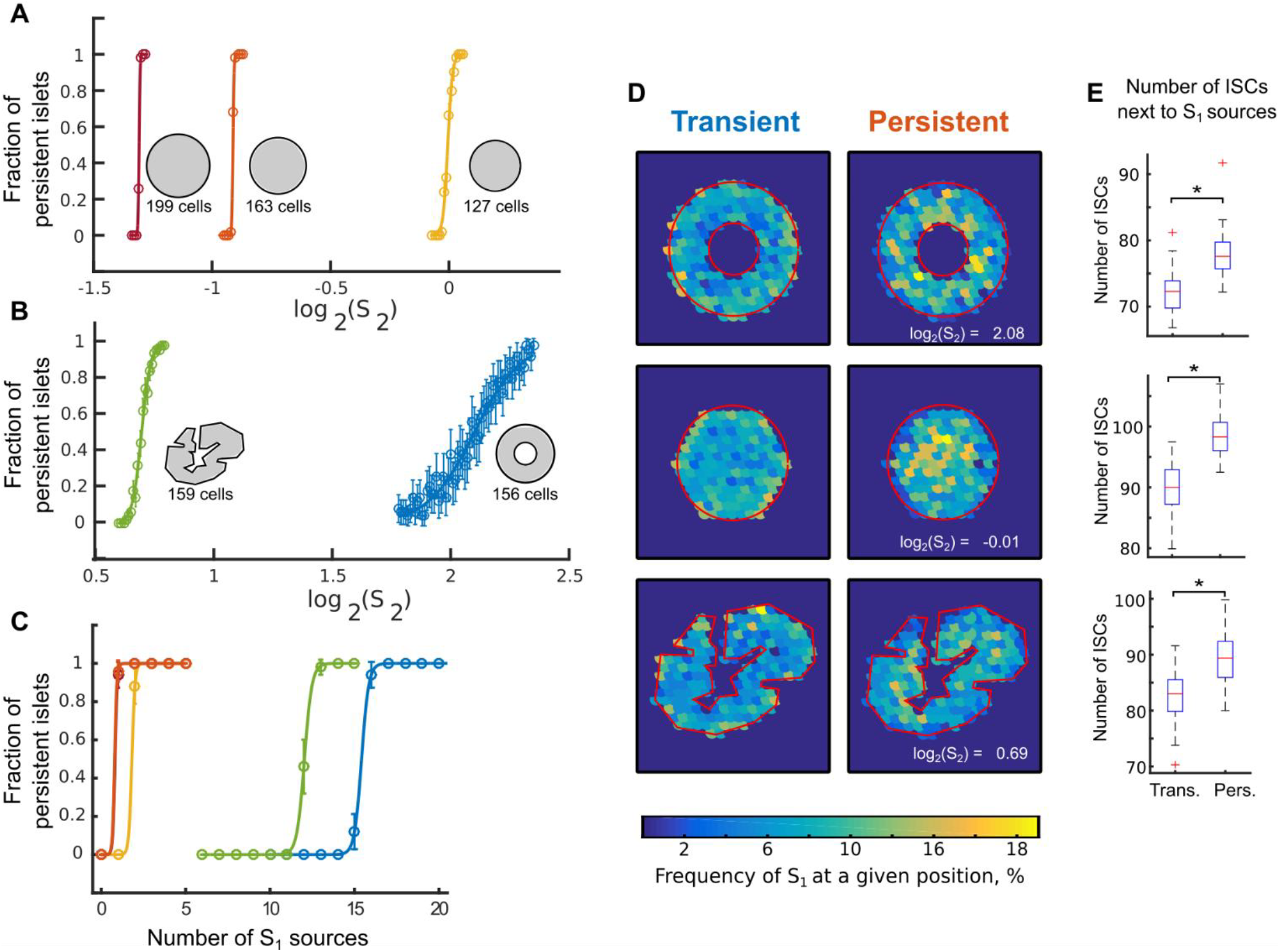
Islet size and shape influence its propensity to lock. **A** Larger islets transition to the persistent mode at lower *S*_*2*_. **B** Islets with few or no ISCs in the core sustain high values of *S*_*2*_ without locking. The islet to the left is a cross-section of a human islet configuration experimentally reported in [18]. The results are shown for *S*_*1*_ = 25 hr^-1^ at each source, with number of *S*_*1*_ sources ns=10. **C** Large core-mantled shaped islets are also more prone to locking when exposed to increasing *S*_*1*_. *S*_*1*_ at each source is 25 hr^-1^, log_2_(*S*_*2*_)=0.5. **D** The sources of *S*_*1*_ positioned deep in the bulk of the ISCs increase islet propensity to lock. The frequency of *S*_*1*_ at a given position is color coded in islets that either respond transiently (left) or persistently (right). Results are sampled over 200 simulations where the position of 10 sources of *S*_*1*_ where chosen randomly among ISCs while other parameters were kept constant (*S*_*1*_ =25 hr^-1^). **E** The density of ISCs in the vicinity of *S*_*1*_ sources is estimated by the average number of ISCs within diffusion length 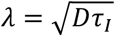 from *S*_*1*_. For all shapes, the number of ISCs next to *S*_*1*_ sources is significantly smaller in transiently responding islets. (Kolmogorov-Smirnov test, comparing the distributions for each islet fate, see Methods). All simulations were run with P=1400.

To further quantify how the transition depends on the shape of the islet, we have implemented an experimentally reported configuration from [18] (Figure 5B, left). Despite relatively large number of cells (159 cells), this configuration is more robust and can sustain higher levels of 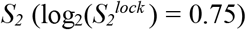 without locking compared to smaller islets with core-mantle configuration (127 cells, 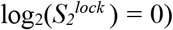). Consistent with this result, islets with ISCs distributed in a donut-shape, transit to the locked state at even higher *S*_*2*_. The results are qualitatively the same if we fix levels of *S*_*2*_ and increase *S*_*1*_, Figure 5C. If we assume both signals are constant throughout the pancreas, larger islets would be at higher risk of entering a self-sustained inflammatory response, simply because the critical *S*_*2*_ level is lower, consistent with observations in T2D patients and murine models [23]. In our simulations, a greater mixture of endocrine cell types or capillaries that can act as sinks can contain this threat.

For each islet shape there is a parameter range where transiently and persistently responding islets coexist. This is not the result of *S*_*1*_ and *S*_*2*_ but rather the internal geometry of sources in the islet. The position of the external sources is random and is the only source of stochasticity in our simulations. Interestingly the region of coexistence is much broader for the human- and donut-shaped islet, indicating that the position of external sources within these islet types have a greater impact on islet fate. Voids, corresponding to non-ISCs or capillaries, could be a factor that increases fate diversity, since a central sink can create a delay in the excitation between ISCs at different ends of the islet. In compact islets, the excitation will spread more or less uniformly, because we use a high diffusion constant compared with islet size.

To investigate why there is a coexistence of modes, we have quantified if some positions of *S*_*1*_ are more likely to induce persistent modes while others transient. In Figure 5D we show the frequency of source positions in transient and persistent modes. We find that the local density of ISC around the sources are higher in persistent compared to transient modes. This is quantified in the box plot of Figure 5E, where there are significant differences in the average number of ISC neighbors within the diffusion length, 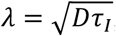, between islets of different fates. Hence an inflammatory response located at the center of an islet is more critical than peripheral cell defects or a response close to capillaries, where the cytokines can be cleared from the islet.

## DISCUSSION

We investigated the possible dynamical modes that can be generated by IL-1 regulatory network in pancreatic islets in response to pro-inflammatory, *S*_*1*_, and metabolic, *S*_*2*_, cues. Notably, we find that there are only two – *transient* and *persistent* – dynamical modes. Weak cues result in *transient* upregulation of IL-1, while strong cues push the system into a state where IL-1 is *persistently* upregulated, even when the initial cues are removed. These modes are encoded for by the fast positive IL-1 feedback and slow negative feedbacks on NF-κB. Islets respond transiently when the strengths of these feedbacks are balanced and persistently when positive feedback is dominating. When the negative feedback is dominating, the response will be diminished or completely ablated. The progression of T2D can be seen as imbalanced feedbacks, whereas beneficial effects of antagonists including IL-1R antagonists [39] to pro-inflammatory cytokines and chemokines, can be thought of as a tool to restore the balance (see Figure 6).

**Figure 6:**
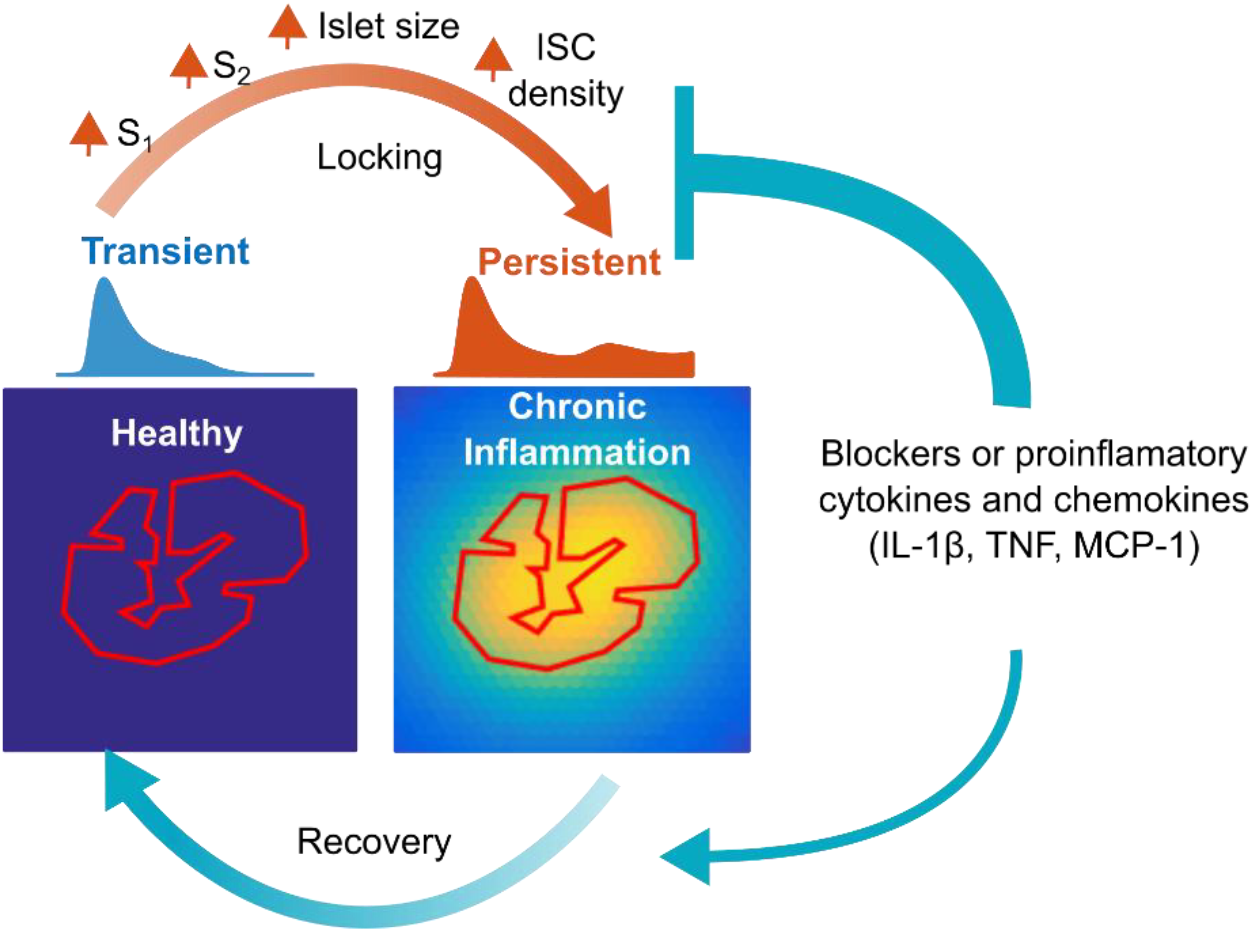
Summary of the results and model predictions. The combination of the negative and positive feedbacks regulating the NF-κB response results in two qualitatively different islet fates. When briefly exposed to pro-inflammatory stimuli, some islets will upregulate IL-1β transiently while others persistently. The increase in the proinflammatory (*S*_*1*_) or metabolic (*S*_*2*_) cues, size of the islets or density of IL-1-Secreting Cells (ISC, β-cells or macrophages) increases the likelihood for the islet to respond persistently. Blockers of pro-inflammatory cytokines and chemokines weaken the positive feedback loop and offset the transition into persistent mode (flat-headed arrow). Higher doses can reverse the transition; however concomitant reduction in *S*_*2*_ is required for the long-term effect upon treatment withdrawal.

### Offsetting and reversing the transition by IL-1 receptor antagonists

While it is often mentioned that the beneficial effect of IL-1 antagonism is derived from “*interruption of vicious cycles of auto-inflammatory induction”* [39], our results introduces a note of caution against complete interruption of the local auto-inflammatory positive feedback, as it is likely to be essential to mount a protective transient response. Instead of “interrupting” the IL-1 positive feedback, the aim should rather be at “restoring” the balance between the two feedbacks (which is likely what happens at the administered doses of IL-1 antagonists). We modeled this effect by weakening the positive feedback, but an alternative way of restoring the balance, would be to strengthen negative feedbacks. In that respect, the recent results with lysine deacetylase inhibitor Givinostat [40] pose an interesting question of how hyperacetylation of the p65 subunit, a component in the pathway that should affect strengths of both feedbacks, alters the balance between them.

Our simulations with IL-1Ra suggest that disrupting the autoinflammatory positive feedback *at the single islet level* is necessary, but not sufficient to explain the observed long-term improvement in T2D patients. It can however explain the long-lasting effect when combined with a decrease in inflammasome activity, the *S*_*2*_-value. In single islets this decrease may happen either locally, for example through the pathway of pro-inflammatory cytokines→ER-stress →>inflammasome [3, 41, 42] or through a global, systemic effect, where improved β-cell function leads to normalized glycaemia and decrease in inflammasome activity.

Our results are thus consistent with the recent in-silico models of the therapeutic effect of IL-1Ra focusing on whole-body effects by Zhao et al. [43] and Palmer et al. [44]. Both models implement the dual effect of IL-1β on proliferation and apoptosis and predict how treatment affects β-cell mass. In an extensive disease progression model, Palmer et al. is able to reproduce sustained improvements in insulin secretion after a *transient treatment* with IL-1RA and conclude that this is mainly due to an improvement in β-cell function rather than β-cell mass [44]. In line with this, Zhao et al. find that treatment with IL-1Ra *must be transient* to see disease improvement as the treatment also inhibits the proliferation of β-cells. This may explain why IL-1Ra or IL-1β antibody therapy for 9-12 months failed to improve beta-cell function in T1D patients [29]

It is tempting to consider that IL-1Ra may be yet another component of the “combined” slow negative feedback of the IL-1β->IKK->NF-κB pathway. There are NF-κB binding sites upstream of the IL-1Ra promoter [45]; is reported to be upregulated by IL-1β in endometrial stromal cells [46]; the expression of IL1Ra follows expression of IL-1β when induced by LPS in Schwann cells [47], and both are upregulated in response to leptin [48] in β-cells. Thus, as is the case with our combined negative regulator, *R*, IL-1Ra may be upregulated transiently because of transient IL-1β activation when the positive and negative feedbacks are balanced.

### Large and dense islets are more likely to transit into persistent mode

Our model predicts that in the population of islets exposed to the same conditions both transiently and persistently responding islets will be observed. This co-existence is a result of geometrical differences in islet structures or in spatial configurations of initial pro-inflammatory cues.

*Large* and *dense* islets, where many of the ISCs are next to each other, are more likely to transit into the *persistent* mode. Due to extracellular diffusion, ISCs surrounded by non-ISC are exposed to lower IL-1 than the cumulative IL-1 secreted by several neighboring ISCs. Thus effectively higher density of ISCs corresponds to a stronger positive feedback experienced by individual cells. Therefore, large and dense islets will be biased to respond persistently. Notably, Striegel et al. [49] found that β-cells are more proximal to each other in islets of T2D patients, which suggests that all the other parameters equal, islets of T2D patients are more likely to transit into persistent modes.

These results may unify the experimental observations by Ehses et al. [15] and Youm et al. [24] and provide an explanation to a hitherto unanswered question of how size of the islets may relate to their inflammatory state and eventual death in T2D [23]. They also may suggest an evolutionary perspective on limiting the size of the islets by distribution into 1 x 10^6^ separate micro-organs dispersed in the exocrine pancreas: While it has been proposed that the size of the islets may be optimized for synchronization in Ca^2+^ bursts and glucose induced insulin release [50], our results suggest that clustering β-cells in physically separated small islets may serve an additional purpose of reducing inflammation and isolating it to single islets.

The coupled fast positive and slow negative feedbacks classify IL-1β regulation as excitable media phenomena[51]. It is well established that information processing or “chemical computing” can be performed by certain spatial configurations of excitable and non-excitable units [52]. Our results, showing that the transition from protective (transient) to deleterious (persistent) modes strongly depends on islet size and the geometrical configurations of excitable ISCs, open for an exciting possibility that the spatial organization of the islets may reflect “cytokine computing” at the islet level.

### Outlook and suggestions for experimental validations

Our results call for quantitative experiments at the single islet level. While technically challenging, it now becomes conceivable to sample cytokine secreted by single islets using microfluidic devices [53]. However, antibody staining and RNA FISH to detect levels of IL-1 protein and mRNA in single pancreatic islets can also validate a number of our predictions. Our model will be validated if with increasing doses of the cues, there will be two subpopulations of islets, one responding transiently and another persistently, and with a bias towards larger islets in the persistent fraction.

Our approach and results are likely to apply beyond IL-1β in pancreatic islets, for example to other pro-inflammatory cytokines (e.g. TNF, IL-6, MCP-1)), where it is known that both positive and negative feedbacks are at the core of the regulation.

## METHODS

### Parameter and variable rescaling

We present the equations in rescaled form (see supplementary for the equations with non-scaled parameters and variables). The variables *I* and *I’* are rescaled with respect to the activation threshold *K*_*I*_, and *N* and *R* are rescaled with respect to total number of NF-κB proteins *N*_*tot*_. The model only has *S*_*1*_ and *S*_*2*_ as free parameters, where *S*_*1*_ is rescaled with respect to *K*_*I*_.

We hand-fitted *k*_*IN*_, *k*_*RN*_, *k*_*NR*_ and *τ*_R_ to fit the time-scale of the first peak in NF-κB oscillations [30]. The timing of the first peak in the model has a small delay and is a bit wider than the peak of actual NF-κB oscillations. This is because several inhibitors are collapsed into the *R*-variable, which also removes the secondary NF-κB oscillations normally observed.

**Table 1:**
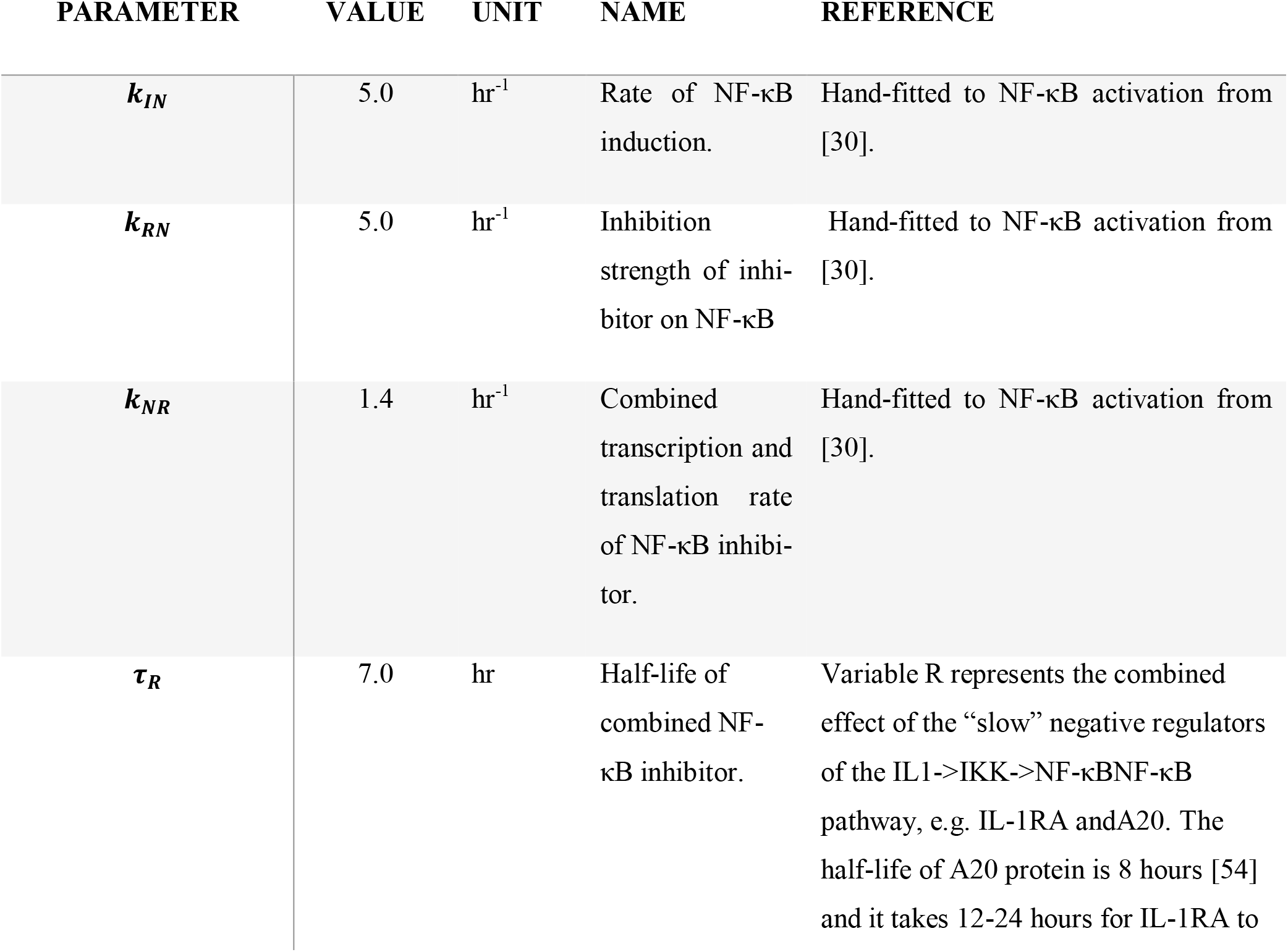
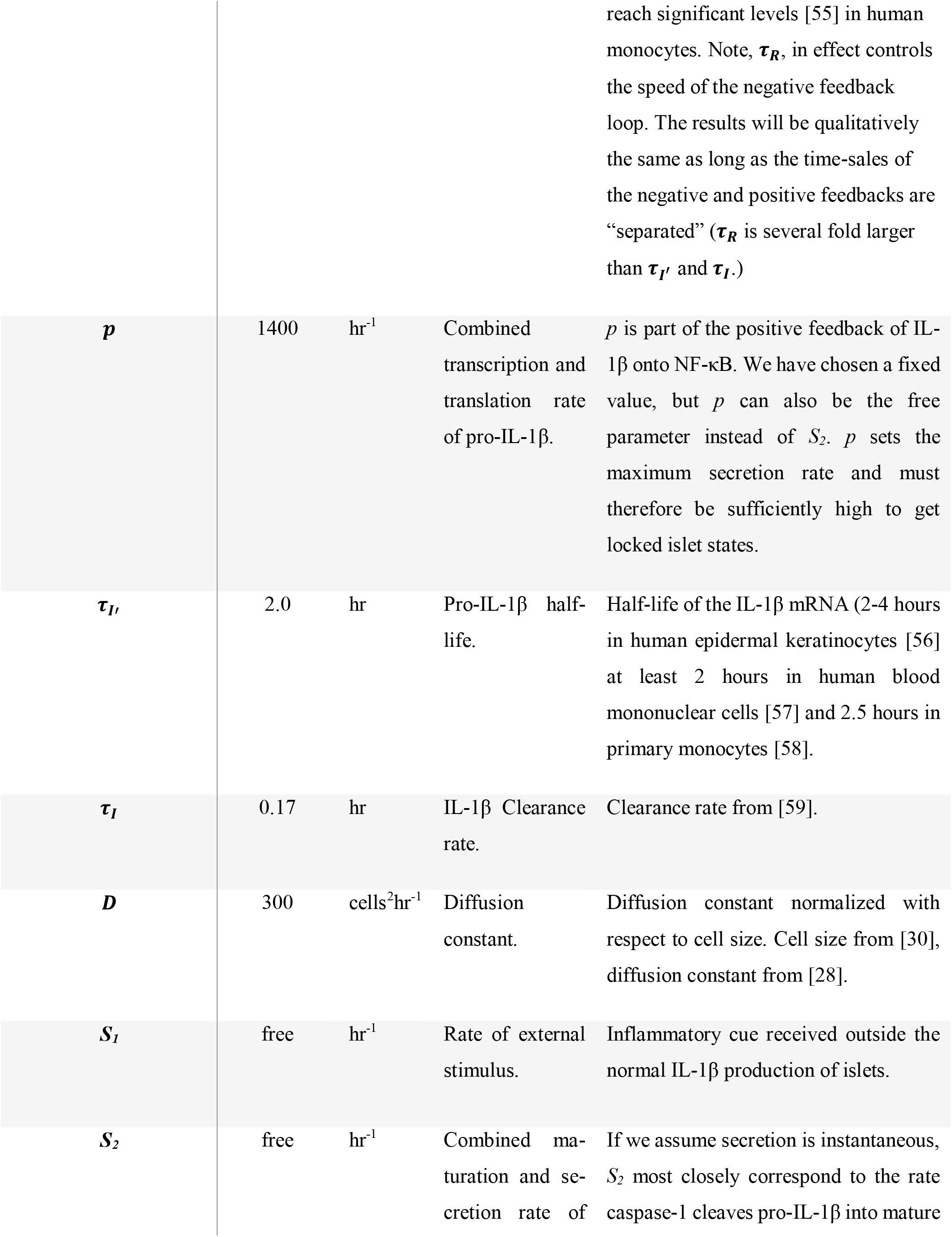
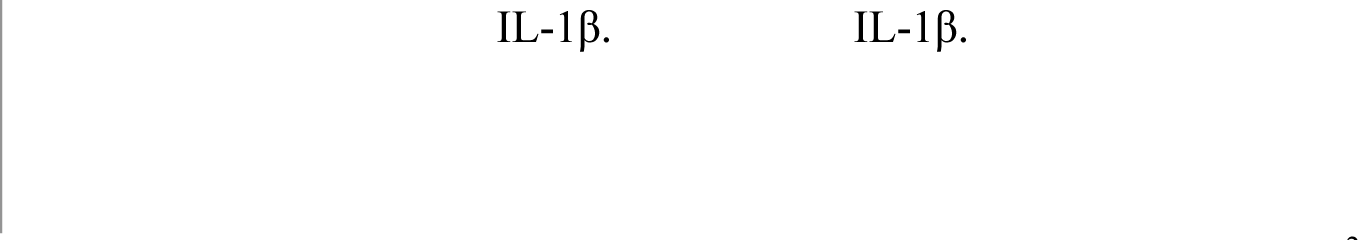
Model parameters.

The diffusion constant is in units of cell size. We used a cell size of 15µm [18] and D = 900 µm^2^/min [28] the diffusion on a grid of cells is D ≈ 300 cells^2^/hr. Note, IL-1β can be exported in a variety of ways; exocytosis, microvesicles, by the use of membrane transporters and upon lysis [60]. We have not modeled the export step explicitly, but it can introduce a further delay between NF-κB-activation and mature extracellular IL-1β.

### Numerical integration and analyses

The ordinary differential equations were solved numerically in C++ and the analysis in Matlab. We organized the cells on a hexagonal grid to have equal distance to nearest neighbors. Each IL-1 producing cell was programmed as an object with local concentrations of *N, R* and *I’*. We numerically integrated the ordinary differential equations using fourth order Runge-Kutta method and the diffusion and decay of IL-1β with a forward Euler method. We solved the decay of IL-1β method with the Euler method, because the cytokine decays everywhere on the grid. By taking this process out of the IL-1 producing cells, we can limit the number of objects to IL-1 producing cells. Notice that the hexagonal grid calculates the second derivative along three axes. In 2D, diffusion only consists of two derivatives, therefore we multiply the forward Euler by 2/3.

We performed Kolmogorov-Smirnov test using Matlab’s built in function *kstest2*. The test has the benefit that it makes no assumption on the underlying pdf.

Transitions were fitted to a sigmoidal function: 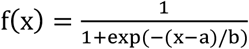, where *a* is the EC_50_-value, where 50% percent of the islets are in the persistent mode.

## ACKNOWLEDGMENTS

This work was supported by the Danish National Research Foundation through the Center for Models of Life and StemPhys.

## SUPPLEMENTARY

### I.a Model: Single Cell Simulations

The model used in this work, is based on the three-variable NF-κB model by P. Yde et al. [25]. We modified the model to increase similarity with the system of pancreatic β-cells, i.e. introducing pro-IL-1β and IL-1β. The fourth variable, pro-IL-1β, was included since glucose affects IL-1β secretion through inflammasome activation, and maturation of caspase-1 which cleaves pro-IL-1β into mature IL-1β [61]. This introduces a new dynamic compared with the old system, since a low signal 2 can lead to a build-up of pro-IL-1β, which was not possible in the old model. In the regime of high signal 2, the translation rate of pro-IL-1β, *p*, can be a limiting factor.

The model takes the following form before rescaling:

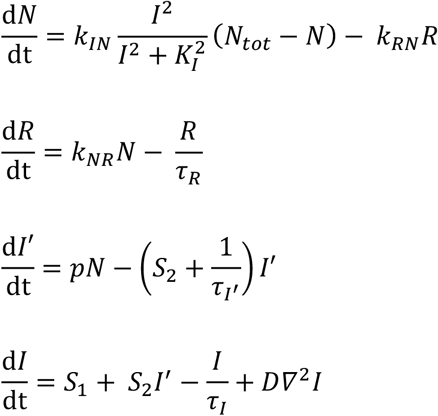

*N* is the nuclear NF-κB, *R* is an effective inhibitor of NF-κB, *I’* is pro-IL-1β and *I* is mature IL-1β. The inactive NF-κB, *(N*_*tot*_ *- N)*, is resident in the cytoplasm but can be activated by IL-1β binding to surface receptors. A Hill- term describes the receptor binding and transportation of NF-κB into the nucleus. The Hill-term, which is non-linear, is the source of bi-stable behavior in the system. NF-κB upregulates both *I’* and *R* linearly and dependent on the *S*_*2*_ parameter, *I’* is transformed into *I* which can diffuse between cells in an auto- and paracrine manner. For simplicity the model assumes the degradation of *R, I’* and *I* is linear as expected by dilution. Active degradation is normally described by Michelis-Menten kinetics but reduces to a linear term in the saturated regime. For simplicity the inactivation of *N* by *R* is modelled as a linear term, but should be modelled by Michelis-Menten kinetics. Because *R* represents several inhibition factors that work at different time-scales, the model does not describe secondary oscillations normally expected by NF-κB. The focus of this model is the transient behavior in response to the initial and sustained stimulus and transitions between qualitative types of response after the initial stimulus is removed.

We rescaled the model to eliminate a few parameters. *N* and *R* are rescaled in terms of *N*_*tot*_ which changes *p* into *p·N*_*tot*_. *I* and *I’* are normalized with respect to the activation threshold *K*_*I*_. This changes *p* and *S*_*1*_ into *p·N*_*tot*_*/K*_*I*_ and *S*_*1*_*/K*_*I*_. The rescaled equations are shown in the main article.

### I.b Model: Critical positive feedback strength

To investigate the strength of the positive feedback consider if pro-IL-1β is in a steady state. The differential equation governing IL-1β can then be rewritten in terms of the steady state value of pro-IL-1β:

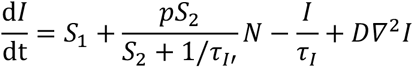

The precursor will not be in a steady state, but the example shows that *p* is a limiting factor in the production of IL-1β. As *S*_*2*_ increases the strength of the total positive feedback also increases, but in a non-linear way. For the human islet and the donut-shaped islet, the system is nearly saturated with respect to *S*_*2*_, and has reached close to its maximum production rate: *p*. To work in the regime of high *S*_*2*_ reactivity, we should choose *S*_*2*_ close to 1/τ_I’_ = ½ hr^-1^.

### I.c Model: Reactivating Inflammatory Response

After an initial inflammatory response the model undergoes a refractory period where it cannot be re-excited, despite returning to the resting state. The length of this period is determined by the amount of inhibitor, *R*, in the system. In the current model re-excitation can occur around 40 hours after the initial response, see Figure S1, but can be reduced by changing the half-life of the inhibitor. The frequency of re-excitations can also increase by changing the strength of the external stimulus, but for too strong a stimulus the islet transits into the locked state. Islets that lock after the first external stimulus, will only have a very limited reaction to additional external stimuli, see Figure S1C.

**Figure S1:**
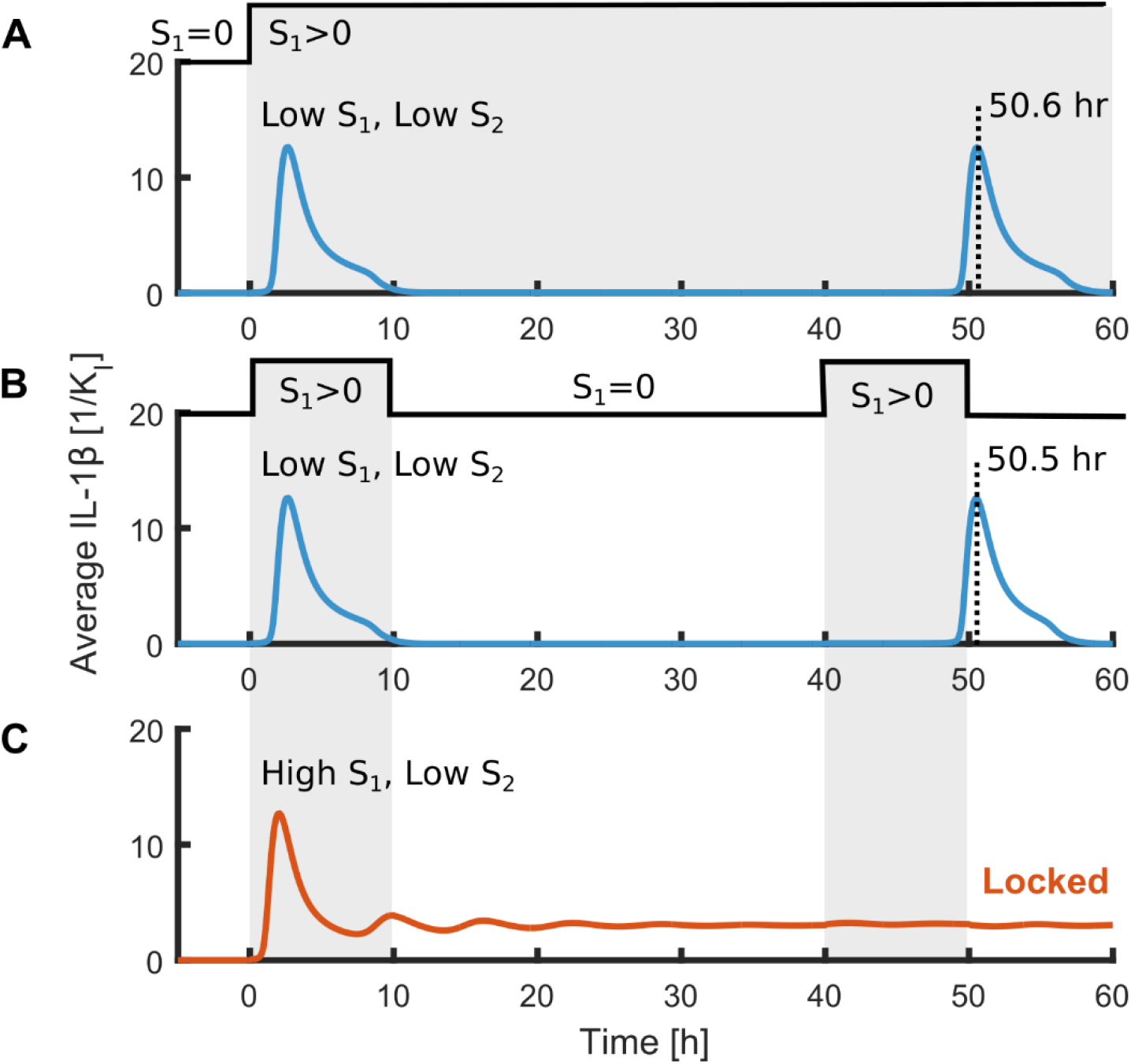
Re-excitation in response to a constant stimulus **A**, and a second pulse of external signal **B**. During the refractory period the system cannot be re-excited due to elevated inhibitor levels. In the case of a pulsing external stimulus the system re-excites slightly earlier, because the external stimulus leads to a slight increase in inhibitor level. If the islet locks after the first stimulus **C**, an additional pulse will have nearly no impact on the level of inflammation.

### I.d Model: Systemic Stimulus

We have considered an initial stimulus as spatially confined sources which produce IL-1β. However, type II diabetics are also subject to systemically increased level of IL-1β, which potentially can trigger a response in the pancreatic islets. In figure S2, we show that there are no qualitative difference between the response from a globally increased basal level of IL-1β and localized sources as shown in figure 2 of the main article. We implemented the basal level by adding a small production term, we also call *S*_*1*_, at each grid point. The source positions is the only source of noise in the model, so by replacing the sources with a basal level of IL-1β, we remove the co-existence of islets with different fate. This would reappear if we considered a distribution of islets sizes or introduced noise elsewhere in the regulatory network.

**Figure S2:**
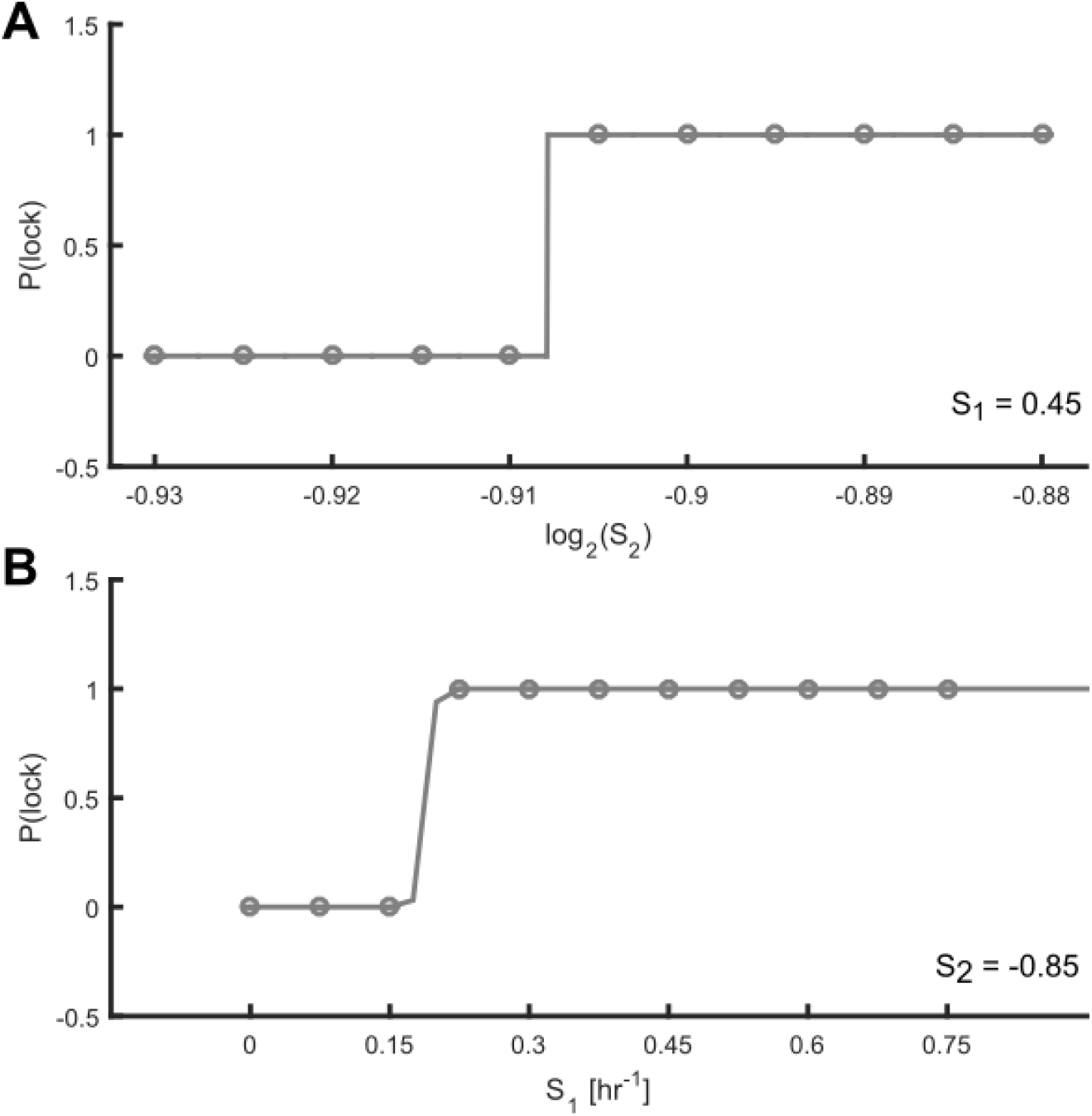
Transition from the transient to the persistent state with a systemically increased basal level of IL-1β. The transition as function of S_2_, **A**, and the basal production rate S_1_, **B**, is abrupt and without a region of co-existence. The overlapping region disappears, as the position of sources were the only source of noise in the model.

## Model: Expanded Parameter Scan

**Figure S3:**
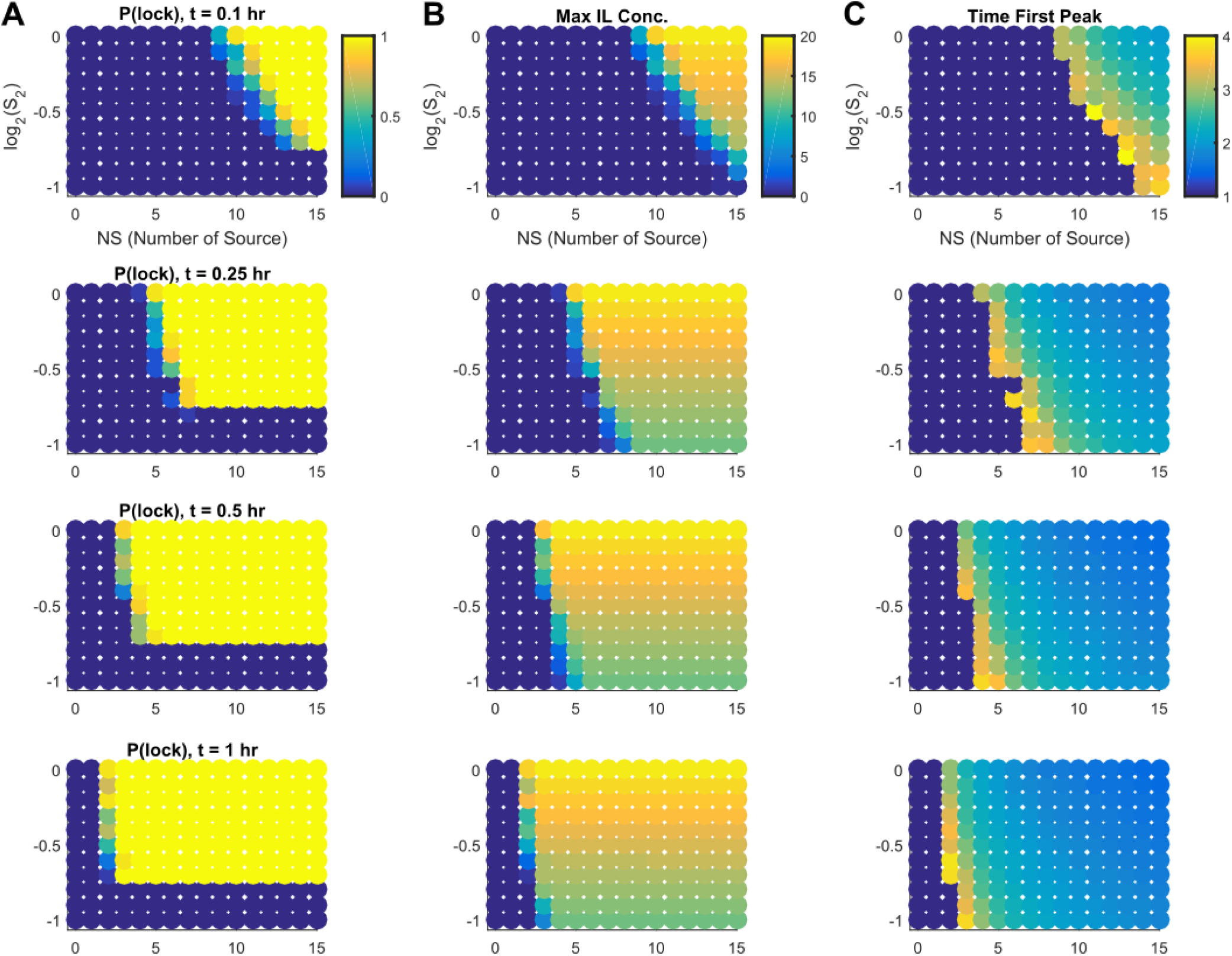
Parameter scan for increasing durations of IL-1β exposure. S_1_ = 25 hr^-1^ per source and each parameter set is the average of 20 simulated islets with random source positions. **A** Fraction of islets in the locked state as function of signal 1 and 2. (I-L) **B** Maximum IL-1 concentration is observed at high S_2_-values. The value is nearly independent of S_1_. **C** The timing of the first peak is mainly dependent on S_1_. The first peak is delayed at the interface between transiently and persistently responding islets, compared to purely persistently responding islets.

**Figure S4:**
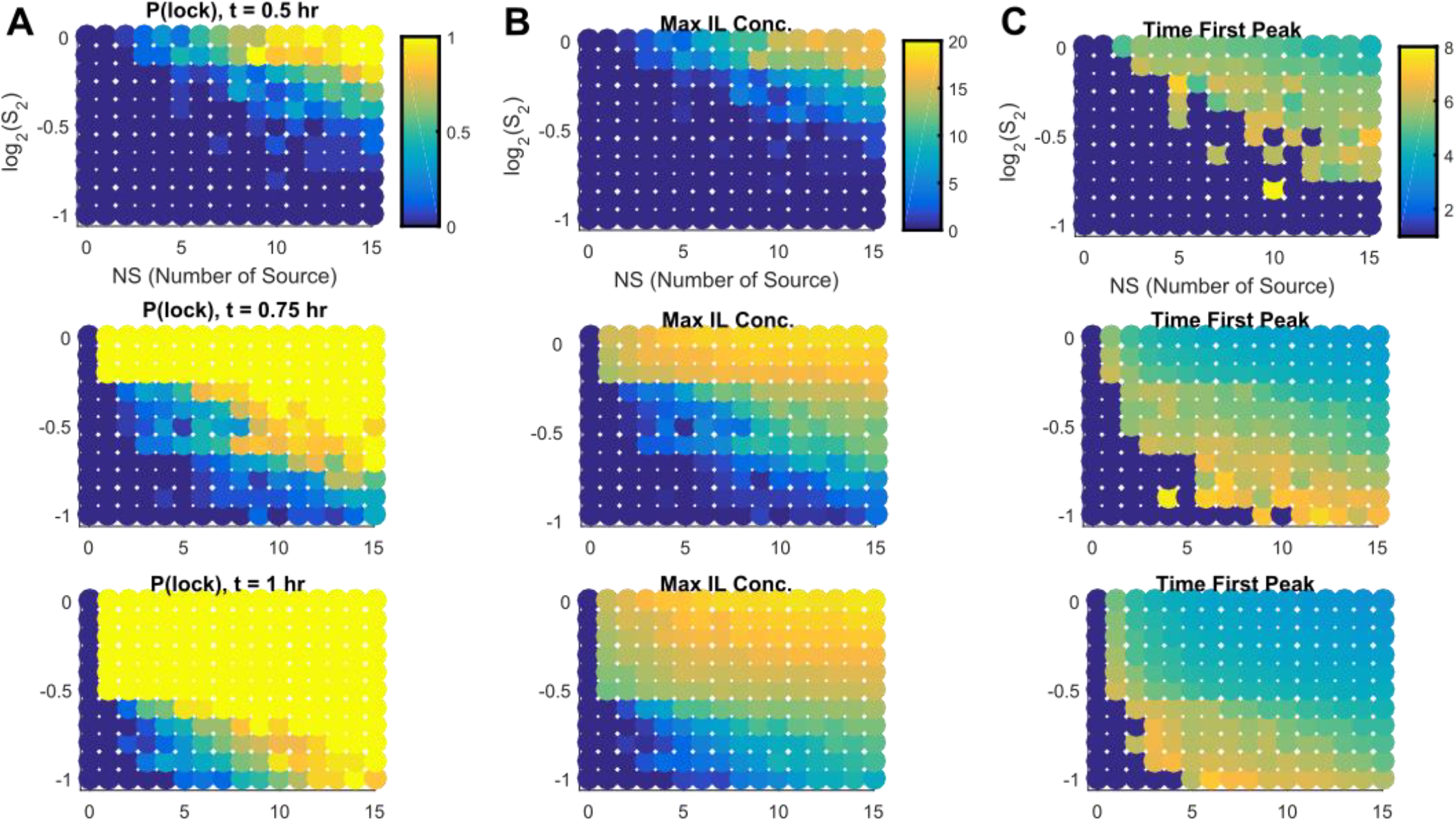
Parameter scan for low diffusion, D = 2 cells^2^hr^-1^. The parameters have been rescaled to get the same qualitative behavior as for high diffusion: S_1_ = 0.5hr^-1^ per source, p = 500 hr^-1^. Averages are over 20 islets with random source positions. **A** Fraction of islets in the locked state as function of signal 1 and 2. (I-L) **B** Maximum IL-1 concentration is observed at high S_2_-values. The value is nearly independent of S_1_. **C** The timing of the first peak is mainly dependent on S_1_. The first peak is delayed at the interface between transiently and persistently responding islets, compared to purely persistently responding islets.

## Analysis

The transition from transiently to persistently responding islets can be fitted with a sigmoidal function. From the sigmoidal function we can extract the EC50-value, which measures where 50% of islets are locked.

**Figure S5:**
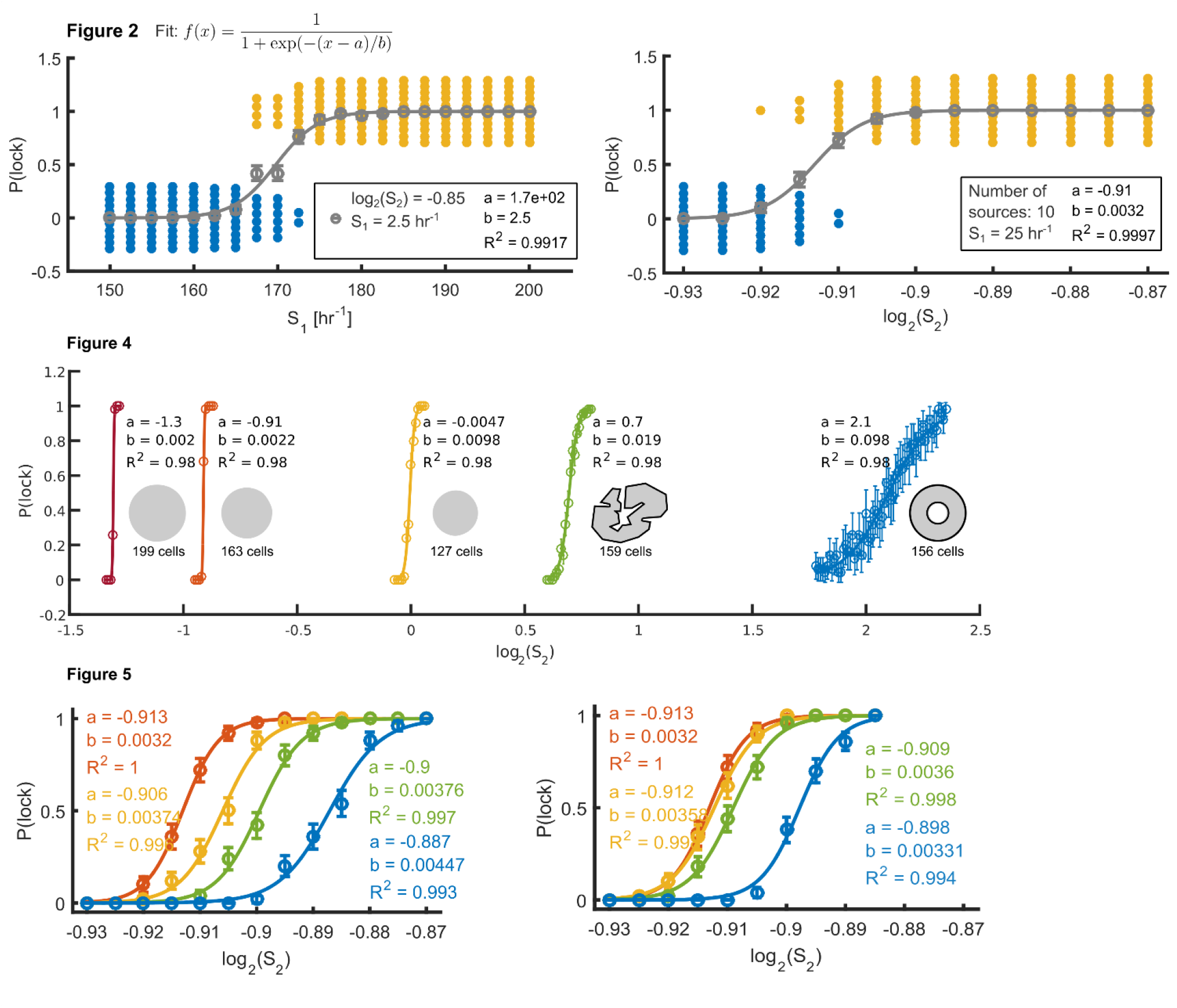
Fit statistics from figures 2, 4 and 5. The transition in lock/persistent probability is fitted to a sigmoidal function, where the fit parameter a is EC_50_, the parameter value, where 50% of islets are in the persistent mode. The fit was done with the built-in fit-function in Matlab.

## Supplemental Movies

- Movies corresponding to islets in **Figure 2: S1:** *LowS1LowS2*, **S2:** *HighS1LowS2*, **S3:** *LowS1HighS2*.
  ∘ *R* = 7 cells, *S*_*1*_ *=*25 hr^-1^, *p* = 900 hr^-1^, *D* = 300 cells^2^hr^-1^
- Movies of large islet: **S4:** *LowS1LowS2*, **S5:** *HighS1LowS2*, **S6:** *LowS1HighS2*
  ∘ *R* = 10 cells, *S*_*1*_ *=*25 hr^-1^, *p* = 900 hr^-1^, *D* = 300 cells^2^hr^-1^
- Movies of low diffusion islets: **S7:** *LowD_LowS1LowS2*, **S8:** *LowD_HighS1LowS2*, **S9:** *LowD_LowS1HighS2*.
  ∘ *R*=10 cells, *S*_*1*_ *=*2.5 hr^-1^, *p* = 500 hr^-1^, *D* = 2 cells^2^hr^-1^
- Movies of low diffusion travelling excitation: **S10**: *movie_one_sink*, **S11**: *movie_two_sinks*.
  ∘ *R*=15 cells, *S*_*1*_ *=*2.5 hr^-1^, *p* = 600 hr^-1^, *D* = 2 cells^2^hr^-1^, sinks: R = 4 cells.

## Notes

### Competing Interest Statement

The authors have declared no competing interest.

